# DensiTree 2: Seeing Trees Through the Forest

**DOI:** 10.1101/012401

**Authors:** Remco R. Bouckaert, Joseph Heled

## Abstract

**Motivation:** Phylogenetic analysis like Bayesian MCMC or bootstrapping result in a collection of trees. Trees are discrete objects and it is generally difficult to get a mental grip on a distributions over trees. Visualisation tools like DensiTree can give good intuition on tree distributions. It works by drawing all trees in the set transparently thus highlighting areas where the tree in the set agrees. In this way, both uncertainty in clade heights and uncertainty in topology can be visualised. In our experience, a vanilla DensiTree can turn out to be misleading in that it shows too much uncertainty due to wrongly ordering taxa or due to unlucky placement of internal nodes.

**Results:** DensiTree is extended to allow visualisation of meta-data associated with branches such as population size and evolutionary rates. Furthermore, geographic locations of taxa can be shown on a map, making it easy to visually check there is some geographic pattern in a phylogeny. Taxa orderings have a large impact on the layout of the tree set, and advances have been made in finding better orderings resulting in significantly more informative visualisations. We also explored various methods for positioning internal nodes, which can improve the quality of the image. Together, these advances make it easier to comprehend distributions over trees.

**Availability:** DensiTree is freely available from http://compevol.auckland.ac.nz/software/.

---

Phylogenetic analysis through Bayesian methods [Bouckaert et al., 2014, Huelsenbeck et al., 2001] or bootstrapping results in tree sets representing a distribution over trees. A popular method of interpreting such a tree set is to construct a summary tree. Such a summary tree can then be visualised (e.g. in FigTree [Rambaut] However, it is hard to represent the uncertainty in the tree’s topology and distinguish it from uncertainty in internal node heights.

Tree sets can be analysed by listing clades and their distributions, but interpretation becomes cumbersome when large number of taxa are involved. Tree networks such as visualised through SplitsTree [Huson et al., 2006] show all possible branches in the trees if supported by sufficient many trees in the set. Multidimensional scaling of tree sets [Hillis et al., 2005] maps every tree in the set on a two dimensional plane, thus grouping trees that are very similar and separating them from trees that differ. It is not possible to read internal node height distributions from a tree network or a multidimensional scaling image.

DensiTree [Bouckaert, 2010] visualises tree sets by drawing all trees in the set transparently in a single image. This way, when trees agree a lot in their topology darker areas appear in the image while when trees in the set differ lighter areas appear. As a result both topological and node height uncertainty becomes visible in a qualitative way.

Here, we explore ways to use the general idea of drawing all instances in the tree set transparently, and extend it to visualising meta data and phylogeography. Further, we address some issues concerning the layout of a DensiTree, which makes them more intuitive.

## 1 Meta Data

In many types of analysis, additional information such as a population size or an evolutionary rate is associated with each branch. There are two ways to visualise such meta data in DensiTree; the line width used for drawing a tree can be used to show the meta data value, or the node position can be used to reflect the meta data value.

When using line widths, the variation of meta data for a certain branch is shown as a blur if there are not too many competing topologies. A block tree is useful for showing the distribution in the dominating topology. Alternatively, a triangle tree can be used, and the top of two coalescing branches adjusted to the bottom of the joining tree branch (see Figure 1, bottom left).

**Figure 1.**
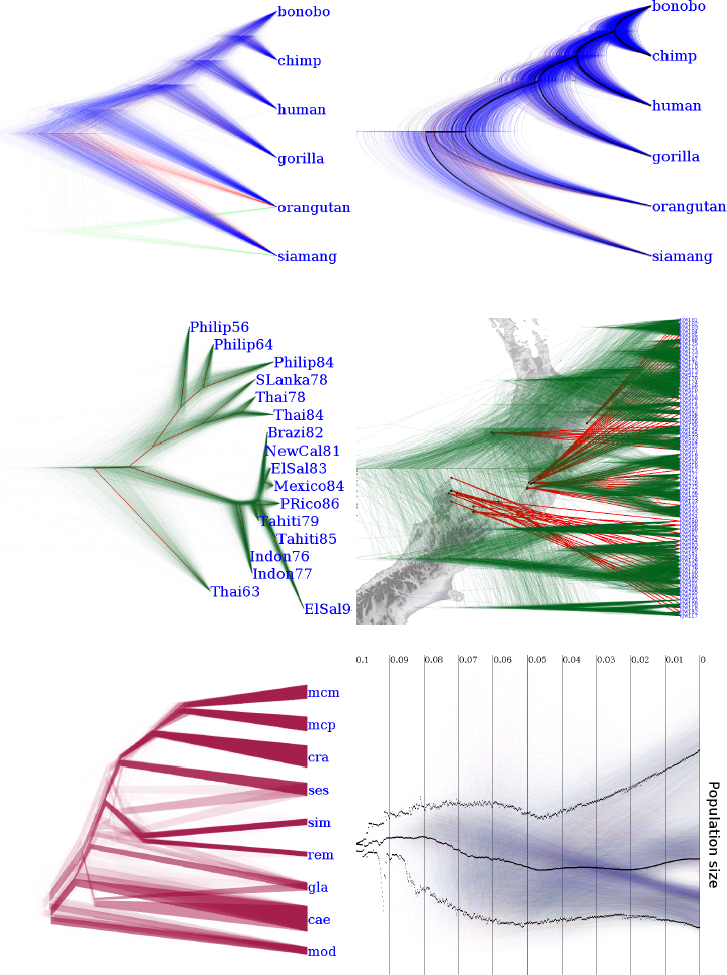
Top left, a centralised DensiTree for great apes. Top right, the same tree set as top left, but with angle correction on the root clade, arced branches and for each topology a mean tree is drawn. Middle left, a star tree with root canal for Dengue virus. Middle right, New Zealand snails and their geographic distribution. Bottom left, Ourisia where banch widths represent population sizes only mean tree for each topology. Bottom right, Ourisia population size with 95% HPD interval and median in black.

When drawing a tree, the height of the internal node is determined by the branch lengths. However, the other coordinate is free to show, say, the population size for the branch. DensiTree allows that coordinate to reflect the meta data value of the branch, drawing every branch of the tree. Alternatively, the sum of all meta data values at a certain height in the tree can be used to draw a single line showing the total population in a tree (see Figure 1, bottom right).

## 2 Phylogeography

Location information of where taxa are sampled is often useful to show together with tree information. DensiTree incorporates known taxa locations (provided in a KML file, which can be exported from Google-earth) by drawing the tree set together with lines connecting taxa with their geographic locations. This allows for a simple visual test whether there is geographical dependence of taxa by finding an ordering of taxa that results in a well defined tree set and where taxa are grouped by location. The middle right plate in Figure 1 shows an example.

## 3 Branch type

Traditionally, trees are drawn as triangle trees or block trees. We explored whether variations on triangle trees would result in clarifying information in a tree set. One issue is that branches that meet come from a different angle, which can give a misleading impression if one of the angles is much less steep than the other. The line at a steeper angle will look much more concentrated, hence appear to have less uncertainty, than lines at a shallower angle.

One solution is to draw branches as part of an ellipse so that the entry angle at the point where branches join is the same (see Figure 1, top right). Alternatively, starting from the child we draw a straight line upwards to the parent, but half the length as drawn in a block tree. Then, we proceed with an arc ending at the parent. Again, angles where lines meet tend to be more similar than when drawing straight lines. Furthermore, this method generally leaves more white space than the partial ellipse method.

## 4 Tip Ordering

DensiTree uses the same tip positions for all trees. As a result, lines must cross when drawing some trees, reducing visual clarity. Finding the order which minimises the overall number of crossings seems hard, but a reasonably fast heuristic can generate a good order in most cases.

For any tree and a pair of taxa define their distance as the size of the clade spanned by their most recent common ancestor, minus 1. So, the distance of two taxa in a cherry (a clade of size 2) is 1. We build a distance matrix *D* for all pairs by averaging distance over all trees. Now, any ordering of the taxa induces a natural distance between pairs given by the absolute difference of taxa positions. A good order would preserve the structure of *D*, keeping pairs with lower distances closer to each other than pairs with larger distances. We define the quality of the ordering by the correlation coefficient between the two distance measures.

Examining all possible *n*! orders is not feasible, and even all 2^*n*−1^ orders compatible with a single tree may take too long. Instead we use a hill climbing approach, selecting a tree through a clustering method or taking the tree with the highest clade support in the set, and flipping the orientation of one internal node at a time, keeping the tree producing the highest quality.

## 5 Internal Node Position

A time tree is visualised by a drawing a path between nodes and their ancestors. The tree provides only one coordinate per node, along the time axis, and the way internal nodes are placed may have substantial visual impact.

By default, DensiTree places an internal node between of its two immediate descendants. This has several undesired consequences. First, the position of a node depends on the sub-tree topology; the root of the same clade can end up in different places for different trees, reducing the intensity of the branch connecting the clade to the tree. Second, long branches are more intense than short branches with the same support. Third, it is hard to separate clades with low support from uncertainty in node heights.

With the new option *star tree* (see Figure 1, middle left) the internal node is placed on the line connecting its parent to the centre of the clade, i.e. between the leftmost and the rightmost taxon. This will centralise the tree, increase the intensity of short branches by drawing shorter and steeper lines, and will compact the area where *fuzzy* nodes are placed. A node is fuzzy if its descendants have low support and are a short branch away. When the nodes are close to each other it appears as if all branches are radiating from the same point, giving a visual clue to the simultaneous radiation at that point.

The new option *centralise* places the internal node between the leftmost and the rightmost taxon (see Figure 1, top plates). Centralising lets the branch keep the same intensity, which now depends only on amount of support for the clade and not on the clade internals. Often, it is clear from the image that a clade with high support has multiple contenders as its child clades. The option *angle correction* automatically detects such clades and puts all of them at the same position as its parent. The result is a tree that looks like more than two arcs coalesce at the same node.

## 6 Technical Details

DensiTree is written in Java 1.6, so no special operating system is required. All techniques discussed above are implemented in DensiTree version 2.01. Drawing takes place using multi-threading for optimal responsiveness and the tree importer is optimised for fast loading of large tree sets. Tree set files can be imported in NEXUS format as produced by BEAST and MrBayes. Images can be exported in bitmap formats (PNG, JPG, BMP) and vector format (SVG). The program is licensed under GPL and source code and a manual [Bouckaert, 2011] are available.

